# Non-cyanobacterial diazotrophs from the Rhizobiales order support marine microalgae in nitrogen-depleted environments

**DOI:** 10.1101/2022.08.25.505241

**Authors:** Udita Chandola, Marinna Gaudin, Camille Trottier, Louis Josselin Lavier Aydat, Eric Manirakiza, Samuel Menicot, Erik Jörg Fischer, Isabelle Louvet, Thomas Lacour, Timothée Chaumier, Atsuko Tanaka, Georg Pohnert, Samuel Chaffron, Leïla Tirichine

## Abstract

**Background:** Non-cyanobacteria diazotrophs (NCDs) were shown to dominate in surface waters shifting the long-held paradigm of cyanobacteria dominance and raising fundamental questions on how these putative heterotrophic bacteria thrive in sunlit oceans. The absence of laboratory cultures of these bacteria significantly limits our ability to understand their behavior in natural environments and, consequently, their contribution to the marine nitrogen cycle.

**Results:** Here, we used a multidisciplinary approach and report an unprecedented finding in the diatom *Phaeodactylum tricornutum* (*Pt*) of NCDs in the phycosphere or the pelagic community sustaining its survival in the absence of bioavailable nitrogen. We sequenced the bacterial metacommunity associated with Pt and assembled several bacterial genomes, identifying multiple NCDs from the Rhizobiales order, including *Bradyrhizobium*, *Mesorhizobium*, *Georhizobium* and *Methylobacterium*. We demonstrated the nitrogen-fixing ability of PtNCDs through in silico identification of nitrogen fixation genes, or by using PCR, acetylene reduction, or 15N incorporation. We showed the wide occurrence of this type of interactions with the isolation of NCDs from other microalgae, their identification in the environment, and their predicted associations with photosynthetic microalgae.

**Conclusions:** Our study underscores the importance of microalgae interactions with NCDs to permit and support nitrogen fixation. This work provides a unique model Pt-NCDs to study the ecology of this interaction advancing our understanding of the key drivers of global marine nitrogen fixation.

## Background

Nearly 80% of Earth’s atmosphere is in the form of molecular nitrogen (N2). Similarly, the main form of nitrogen in the oceans which cover 71% of Earth surface is dissolved dinitrogen. The rest is reactive nitrogen in the form of nitrate, ammonia and dissolved organic compounds, which are very scarce, rendering nitrogen one of the main limiting factors of productivity in most areas of the oceans [1]. Despite its dominance, ironically, N2 is unavailable for use by most organisms. Biological nitrogen fixation which is an energy intensive process is specific to only some prokaryotes and archaea termed diazotrophs. It is catalyzed by the nitrogenase enzyme complex including the nitrogenase reductase encoded by the *nifH* gene and the nitrogenase composed of two pairs of different subunits, *α* and β, respectively encoded by the *nifD* and *nifK* genes [2]. Therefore, eukaryotes are only able to obtain fixed nitrogen through their interactions with diazotrophs [3], although recent evidence has revealed the evolutionary conversion of a cyanobacterium into a nitrogen-fixing organelle within its host, representing the first documented case of nitrogen fixation in a eukaryote [4].

In aquatic habitats, associations between diazotrophic cyanobacteria such as Richelia/Calothrix and various eukaryotes, including several diatom genera such as *Chaetoceros, Rhizosolenia* and *Hemiaulus* known as diatom-diazotroph associations are common and important in oligotrophic habitats of the ocean [5]. Because the nitrogenase complex is sensitive to oxygen, diazotrophs evolved protective nitrogen fixation alternatives that involve conditionally, temporally, or spatially separating oxidative phosphorylation or photosynthesis from nitrogen fixation [6–11]. Cyanobacteria were considered for a very long time as the main diazotrophs in the ocean. However, recent molecular analyses indicated that non-cyanobacterial diazotrophs (NCDs) are also present and active, exhibiting an important presence of diverse *nifH* genes related to non-cyanobacteria, especially to Proteobacteria and Planctomycetes [12–16].

The overwhelming dominance of NCDs raises important questions on how these presumably heterotrophs proteobacteria thrive in the photic zones of the oceans. One emerging hypothesis is that their association with aggregates and larger cellular fractions including phytoplankton, may provide the optimal conditions for nitrogen fixation [16–19]. In our work, we provide evidence for the occurrence of associations between free-living NCDs and several microalgae. More importantly, we identified this interaction in an attractive biological model, the diatom *Phaeodactylum tricornutum (Pt),* opening important and unique opportunities to investigate the molecular mechanisms of diazotrophy in symbiotic interactions in microalgae.

We sequenced the bacterial meta-community associated with *Pt* and identified previously unknown diverse marine diazotrophs of mainly Alphaproteobacteria type, supporting the survival of host cells in the absence of any form of usable nitrogen. Further, we isolated, cultured, sequenced and fully assembled some of the Pt-associated NCDs. Their assembled genomes clustered phylogenetically to NCDs found in the environment and showed environmental co-occurrence patterns with microalgae including diatoms, green algae and haptophytes. Interestingly, most of PtNCDs belong to the Rhizobiales, order known to nodulate legumes. These NCDs include *Bradyrhizobium*, *Rhizobium*, *Mesorhizobium*, *Methylobacterium*, *Phylobacterium* and *Georhizobium* raising important questions into the evolution of symbiotic nitrogen fixation and its terrestrial or marine origins. Our findings differ from a recent study that identified an NCD residing within a diatom as an obligate symbiont with a reduced genome compared to free living NCDs [20]. Our work brings evidence for an apparently prevalent interaction between microalgae and NCDs, that might explain how both partners cope with periods of nitrogen scarcity, and how these NCDs compromise for nitrogen fixation with its high energy cost in fully oxygenated areas of the water column. These new findings will have to be considered in future nitrogen and carbon biogeochemical cycling studies.

## Results

### Identification of non-cyanobacterial diazotrophs interaction with diatoms

In a screening of 10 xenic environmental samples of Pt grown in nitrogen-depleted medium, one accession *P. tricornutum 15* (hereafter *Pt15*), sampled from East China sea, was found to survive in contrast to the axenic control (**Fig. 1A**). This suggests that diatom growth may have been supported by bacterial nitrogen fixation providing a usable source of nitrogen to diatom cells. After 3 months of nitrogen starvation, xenic *Pt15* showed similar growth to non-starved cells upon transfer to nitrate replete medium. This indicated that *Pt15* starved cells remained viable and capable of growth under favorable conditions as evidenced by the presence of brown-pigmented cells (**Fig. 1B**). To demonstrate nitrogen fixation, we used the widely applied Acetylene Reduction Assay (ARA) [21] which detected ethylene production in nitrogen-starved xenic *Pt15* cultures. Although the ethylene levels were low, they were notably higher than those in the *E. coli* negative control, where no ethylene production was detected. Xenic *Pt15* cultures under nitrogen-replete conditions (+N) did not show any ethylene production. No ethylene was detected in axenic *Pt15* confirming that the measured ethylene was not endogenous to the diatom cells but originated from bacterial nitrogen fixation **(Table S1**).

**Fig. 1.**
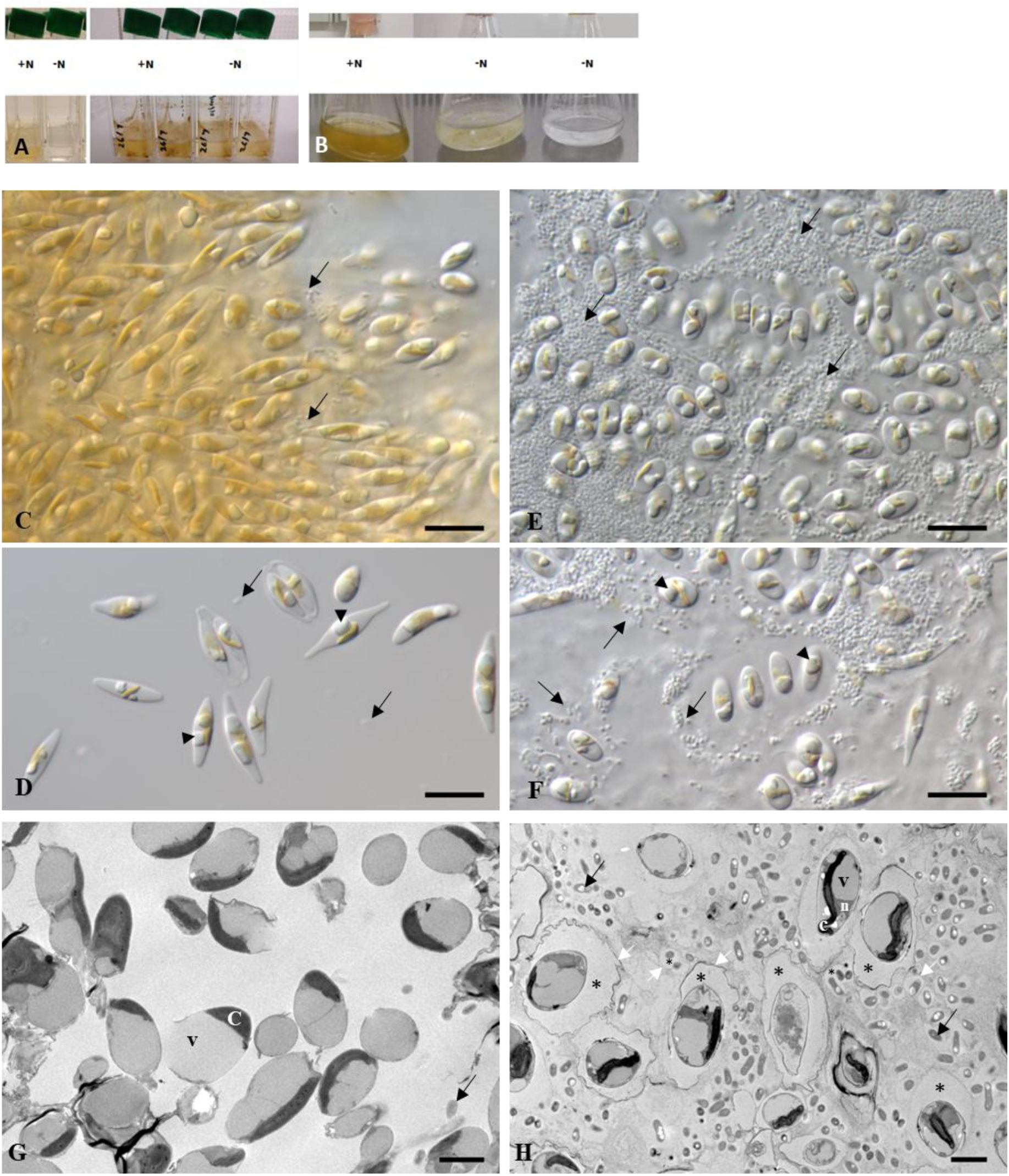
Macro/microscopic phenotype of *P. tricornutum* cells under nitrate starvation. A. An illustration comparing the screening results of two P. tricornutum accessions: xenic Pt4 (left), which thrives in +N but does not survive in -N, and the candidate accession, xenic Pt15 (right), shown in duplicates. B, from right to left: axenic Pt15 not surviving in -N (negative control), xenic Pt15 in -N condition and xenic Pt15 transferred to +N medium after 3 months in -N. C-H, micrographs of P. tricornutum xenic cultures on plates. Cells were grown in -N (E, F, H) or +N culture plates (C, D, G). Living cells were observed by DIC (C-F), and fixed cells by transmission electron microscopy (G, H). D, Diatom cells grown in +N possessed yellow-brown color of photosynthetic pigments. Mainly in aggregated cells of fusiform and oval shapes, some bacteria were observed (arrows). D, Single or multiple whitish droplets, oil bodies, were developed next to chloroplasts (arrowheads). E, F, Diatom cells formed aggregates with a significant number of surrounding bacteria (arrows) in the -N culture, compared to the +N condition. F, Diatom cells grown without nitrate also developed oil bodies in the oval cells. G, H, Electron micrographs indicate cross-sectioned cells with or without nitrate. Organelles with high electron density are chloroplast, and major spaces were occupied by oil bodies or vacuoles, which showed low electron densities. Number of bacteria cells (arrows) were obviously larger in -N (H) compared to +N culture (G). H, Interestingly, both Pt cells and some bacteria are encased in sacs surrounded with a likely EPS made membrane (white arrows) which showed darker electron density around diatom cells compared to bacteria. Cells in 1G and 1H were stained with TI blue stain, which binds to negatively charged components such as anionic polysaccharides increasing electron density. A low electron density space (stars) separates Pt cells and bacteria from the membrane. c, chloroplast, v, vacuole, n, nucleus. Scale bars, 10 μm (C-F) and 2 μm (G, H).

To further investigate nitrogen fixation in minus N condition (-N), we measured Particulate Organic Nitrogen quota (PON) in diatom cells and showed a stable level of PON even after 80 days of growth, supporting the maintenance of diatom cells with the amounts of usable nitrogen likely provided by *Pt15* associated bacteria (**Figure S1A, Table S1**). Simultaneously, measured biomass of *Pt15*, indicated a stable cell population suggesting a ground state photosynthetic activity that might support the energy demand of nitrogen fixation (**Figure S1B**). We measured a diatom cell density of 4.3 million cells/ml in the +N condition, whereas in the -N condition, *Pt15* cell density did not exceed 360,000 cells/ml.

It is important to highlight that the growth conditions are extreme, with no nitrate supplied to diatom cells, likely leading to a slowdown in their metabolism to endure the severe conditions. Fixed nitrogen is the sole nitrogen source available to both the diatom and the entire bacterial community, likely resulting in competition for fixed nitrogen. Taken together, our results suggest bacterial nitrogen fixation activity that maintain diatom population cells stable and alive. Light microscopy showed a significant accumulation of bacteria in -N compared to +N condition. This accumulation is not an artifact but is likely a response to nitrate starvation, driven by the formation of biofilm-like clusters of NCDs. These clusters may promote the development of low-oxygen microenvironments favorable for nitrogen fixation (**Fig. 1A, D, E and F**). Bacteria were found surrounding diatom cells in the phycosphere, and not residing inside the cells. Transmission electron microscopy confirmed these observations and revealed a peculiar structure with a higher electron density around diatom cells looking like a pellicle of exopolysaccharides exclusively present in starved cells (**Fig. 1G and H**). Similar structures, although less prominent occurred around some of bacterial cells, either as a single bacterium or a group of several bacteria. The probability that these structures are composed of polysaccharides is supported by the TI blue stain, a well-known dye that has an affinity for staining polysaccharides [22].

To identify species composition of *Pt15* bacterial community in -N and +N conditions, bacterial DNA was extracted and shotgun sequenced revealing a total of 153,4 million reads. Using the GTDB reference database, we identified a total of 2 classes, 12 families, 8 orders and 90 genera. Upon comparison of the two conditions, our analysis identified 90 distinct genera and 3,224 bacterial species in the -N condition, whereas only 18 genera and 1,540 species were detected in the +N condition. At class level, Alphaproteobacteria represented 98% ± 0,8 SD in +N and 99% ± 0,5 SD in -N, based on relative abundance estimations, while *Gammaproteobacteria* were detected with a relative abundance below 1% (**Fig. 2A**). At family level, *Sphingomonadaceae* and *Rhizobiaceae*, both known to contain diazotrophic species, dominated (**Fig. 2C**). Attached bacteria may be underrepresented due to the filtration process used to separate diatom cells from bacteria prior to DNA extraction. However, to maximize the recovery of both attached and free-living bacteria, diatom cells were vigorously vortexed multiple times before filtration.

**Fig. 2.**
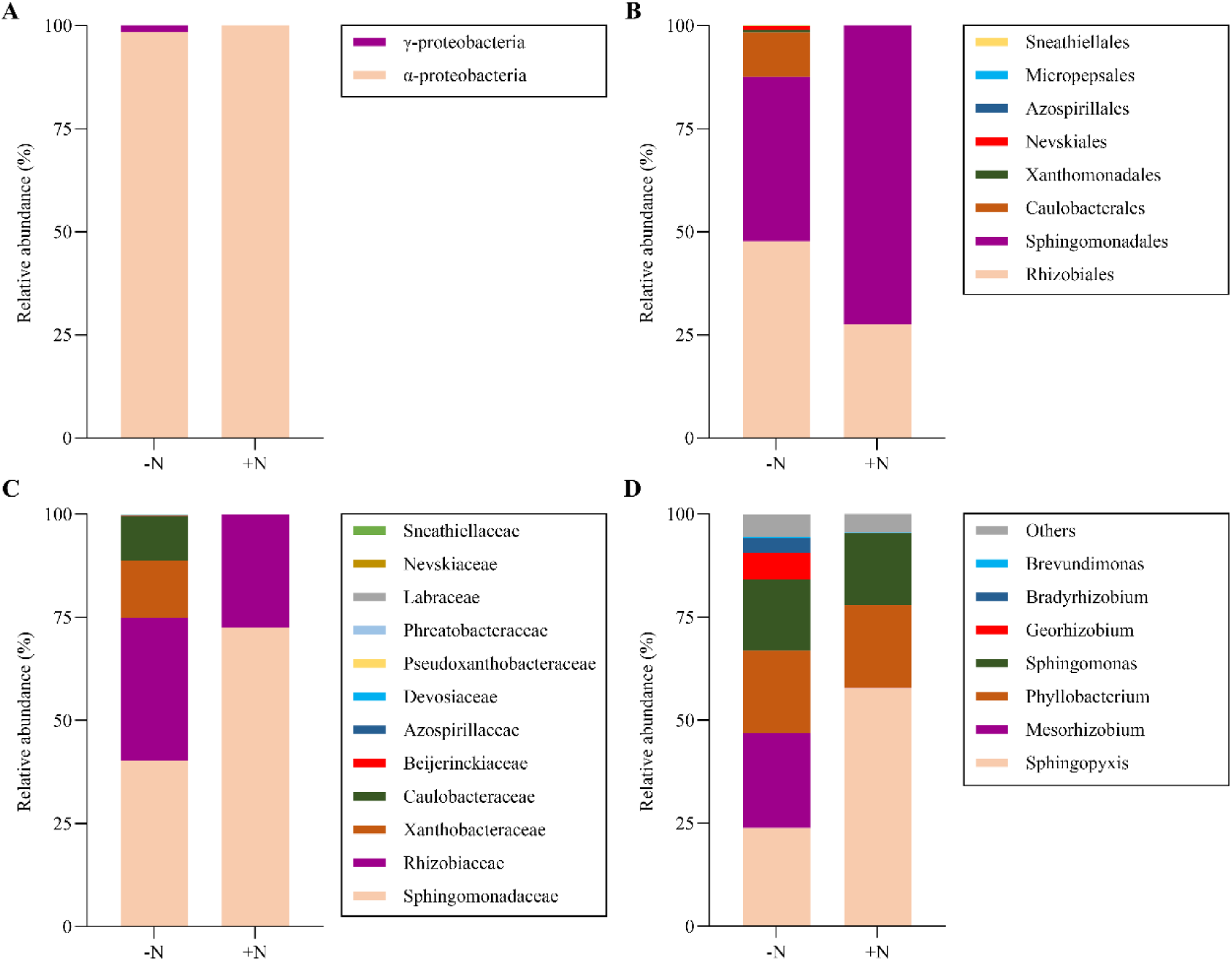
Comparative relative abundance estimation of *Pt15* associated bacteria. Relative abundance of bacterial community is shown as A, class, B, order, C; family and D, genus (7 most abundant) levels between -N and +N conditions. Taxonomic assignation of metagenomic community associated to *P. tricornutum* in nitrate-deplete and nitrate-replete conditions was done using Anvio version 7.2 standard recommended pipeline using GTDB (The Genome Taxonomy Database) for assigning taxa with the in-built HMM of 22 single copy core genes [23].

At genus level, among the 90 distinct taxa identified, 73 genera were only detected in -N with no reads in +N (**Table S2**). At the genus level, the microbiome composition of the community showed a dominance of *Sphyngopixys* in +N, while -N showed a significant enrichment in *Mesorhizobium sp., Bradyrhizobium sp*., *Georhizobium sp*., and *Brevundimonas sp*. These genera are recognized as nitrogen-fixing taxa in terrestrial environments, either existing as free-living organisms and/or forming symbiotic relationships with legumes (**Fig. 2D**). Compared to +N, *Sphyngopixys* abundance decreased in -N although its dominance over the other taxa persisted under this condition. A total of 5 *nifH* genes were detected and assigned to 4 different MAGs (MAG21-Bradyrhizobium: 2 *nifH*, MAG8-Bradyrhizobium: 1 *nifH*, MAG4-Georhizobium: 1 *nifH*, and MAG5-Sphingopyxis: 1 *nifH*) from the -N bacterial community. No *nifH* genes were detected in the +N condition. Interestingly, several presumably non-nitrogen fixing species (67 out of 73) were significantly enriched in -N (**Table S2**), suggesting a role of these species in direct or indirect interactions within the bacterial community and/or with the host. These associated non-diazotrophic bacteria, which are enriched in the absence of bioavailable nitrogen, may contribute to the interaction through various mechanisms. We hypothesize that they could modulate competition among diazotrophs interacting with the host, produce compounds that promote microalgae growth, and inhibit antagonistic organisms, thereby facilitating the interaction with the host. However, these proposed mechanisms have yet to be demonstrated.

### Isolation/identification of *P. tricornutum* associated non-cyanobacterial diazotrophs

Given the unique nature of the identified interaction involving a model diatom species and NCDs, it was important to isolate the diazotrophic bacteria identified *in silico* from metagenomics (MetaG) data in order to further explore this interaction. We used selective media and took advantage of the species assignation provided by MetaG to target the isolation of the desired bacteria according to literature reports [24–27]. Therefore, different growth conditions (media, temperature) were tested (**Additional File 1**) leading to the isolation of 9 axenic bacterial species (**Table 1**).

**Table 1.** List of isolated bacteria from xenic *Pt15* in -N condition. . NCDs are indicated in bold. NA, not assembled, ND, not determined.

| Genus | MAG No. | Size (Mbp) | No. of contigs | *nifH* PCR identification | *nifH* *in silico* identification | Growth on NFM | Predicted plasmids |
| --- | --- | --- | --- | --- | --- | --- | --- |
| ***Mesorhizobium*** | **MAG3** | 6.85 | 75 | No | Yes | Yes | Yes |
| ***Bradyrhizobium sp1\**** | **MAG21** | 8.42 | 5 | Yes | Yes | Yes | No |
| *Sphingopyxis sp2* | MAG7 | 5.00 | 23 | ND | No | ND | Yes |
| *Rhizobium sp.* | NA | ND | ND | ND | ND | Yes | ND |
| *Paracoccus sp.* | NA | ND | ND | ND | ND | Yes | ND |
| *Methylobacterium\** | NA | ND | ND | Yes | Yes | Yes | ND |
| *Arthrobacter* | MAG14 | 3.46 | 12 | No | No | ND | ND |
| *Pseudoarthrobacter* | NA | ND | ND | No | ND | No | ND |
| *Micrococcaceae* | NA | ND | ND | No | ND | No | ND |

NCDs are indicated in bold. NA, not assembled, ND, not determined. \* 15N incorporation was used for nitrogen fixation detection

Bacterial colonies were identified using either full length 16S rRNA gene or two hypervariable regions of the 16S (V1V2, V3V4). The presence of *nifH* was assayed using degenerate primers (**Figure S2, A and B**). Six bacteria were identified as nitrogen fixers based on a combination of *nifH* PCR amplification and/or *nifH in silico* detection along with observation of growth on nitrogen free medium for those that were isolated (**Table 1 and Table S3**). The whole set of *nif* genes including the *nifHDK* operon were identified in *Bradyrhizobium* MAG21, the only bacterium that was sequenced as a single species, while the other MAGs were assembled from MetaG data. Among the nine isolated bacteria, some species were not detected in metagenomics sequences likely due to its low abundance, emphasizing the importance of combining sequencing and isolation/culturing approaches (**Table 1**). Due to the use of selective culturing media for bacterial species isolation, we identified species that were not necessarily present in the in-silico analysis of the metacommunity, which was grown under a single condition.

To provide stronger support for nitrogen fixation, we monitored ^15^N2 gas-incorporation using mass spectrometry. This allowed a direct measure of nitrogen fixation in the two first isolated bacteria, *Bradyrhizobium* MAG21 and *Methylobacterium*. The labeling of various amino acids indicated substantial ^15^N2 gas assimilation in both bacteria as well as the positive controls *Pseudomonas stutzeri* and *Azotobacter vinalendii*, while the negative controls, (the axenic *Pt15* and *Escherichia coli*) did not show any nitrogen fixation (**Table S1**). Our high-resolution LC-MS approach precisely detects the label at the molecular level through isotope-resolved ions, enabling structure-resolved detection of labeled compounds. This allows for more accurate identification of 15N2 incorporation patterns, which would be harder to resolve with bulk detection methods. This detection unequivocally establishes the nitrogen-fixing capability of the *Pt15*-associated bacteria, namely *Bradyrhizobium* and *Methylobacterium*. We could not observe ^15^N2 incorporation in xenic *Pt15* cells, which can be attributed to their lower bacterial population density when compared to the lab cultured high biomass of *Bradyrhizobium* and *Methylobacterium*, as well as the competition with other numerous bacteria (∼ 100 species) for fixed nitrogen. To verify that the ability of *Pt15* cells to uptake bioavailable nitrogen was not altered, we cultured xenic *Pt15* in artificial seawater containing L-valine labeled with ^15^N, both in the presence and absence of nitrate. The analysis of endometabolomes using LC-MS distinctly revealed the incorporation of labeled valine in xenic *Pt15* when nitrate levels were at 0 µM (**Table S1**). However, no such incorporation was observed in diatom cells when nitrate was present in the medium (545 µM). This suggests that *Pt15* is able to take up nitrogen in form of amino acids from the associated NCDs, if available nitrogen is otherwise limited.

### Diazotrophic MAGs are diverse and affiliated to environmentally sampled diazotrophs of Alphaproteobacteria type

To gain insights into *Pt15* associated diazotrophs and their phylogenetic relationship with known taxa, we assembled their genomes and recovered several MAGs with completeness exceeding 97% (**Table S3**). To achieve a better identification of the bacterial genetic material, we sought to recover plasmids from non-assembled reads, which are important elements for bacteria survival susceptible to contain nitrogen fixation genes. We used three different plasmid prediction tools and recovered 42 plasmids assigned to different bacterial species with an average length of 97013 bp (**Table S3).** Besides *nifH*, several additional *nif* genes were detected in most of the MAGs, among which two were found in plasmids, *nifU* and *nifS*.

Phylogenomic analysis of *Pt15* MAGs, MAGs of known terrestrial and marine diazotrophs from the MicroScope database [28] and MAGs from *Tara* Oceans data using anvi’o [29] was performed. The genomes from MicroScope were selected by individual phylogenetic analysis of each nine *Pt15* MAG deposited on the platform. Thus, the relatedness on the MAGs represented are to diazotrophic marine (purple), non-diazotrophic marine (white), diazotrophic terrestrial (green) and non-diazotrophic terrestrial (green) bacteria. The phylogeny was built using Anvio platform and in built HMM of 71 scgs (single copy core genes), Bacteria_71. The analysis revealed two clusters of *Pt15* MAGs on the phylogeny, cluster A (blue) and cluster B (red) (**Fig. 3**). Cluster A (*Mesorhizobium*, *Bradyrhizobium* MAGs, *Georhizobium*, *Phylobacterium*, and *Brevundimonas*) was affilated to diazotrophs, while, cluster B (2 *Sphingopyxis* MAGs and *Sphingobium*), was related more to denitrifiers. In both clusters, whenever the related MAGs were not identified as diazotrophs (because of the absence of *nifHDK*, likely due to incomplete assembly), they were found to belong to Rhizobiales, Parvibaculales, Caulobacterales and Sphingomonodales, orders known to contain nitrogen fixing species [30–32]. *Sphinopyxis* MAG5 was identified with *nifH* gene *in silico* and was found to cluster with non- or un-identified diazotrophs. For a clearer depiction of Figure 3, an interactive phylogenetic tree is available on the following link: https://ncdstree.univ-nantes.fr. Altogether, our results including (i) survival of the xenic diatom in the absence of usable forms of nitrogen, (ii) nitrogen fixation demonstrated by ARA and ^15^N-incorporation, (iii) detection of *nifH,* and (iv) the isolation of NCDs from the xenic *Pt15* clustering with known proteobacteria diazotrophs namely rhizobia, indicate that an interaction for nitrogen fixation likely occurs in *Pt15* xenic cultures, supporting the survival of diatoms in -N condition.

**Fig. 3.**
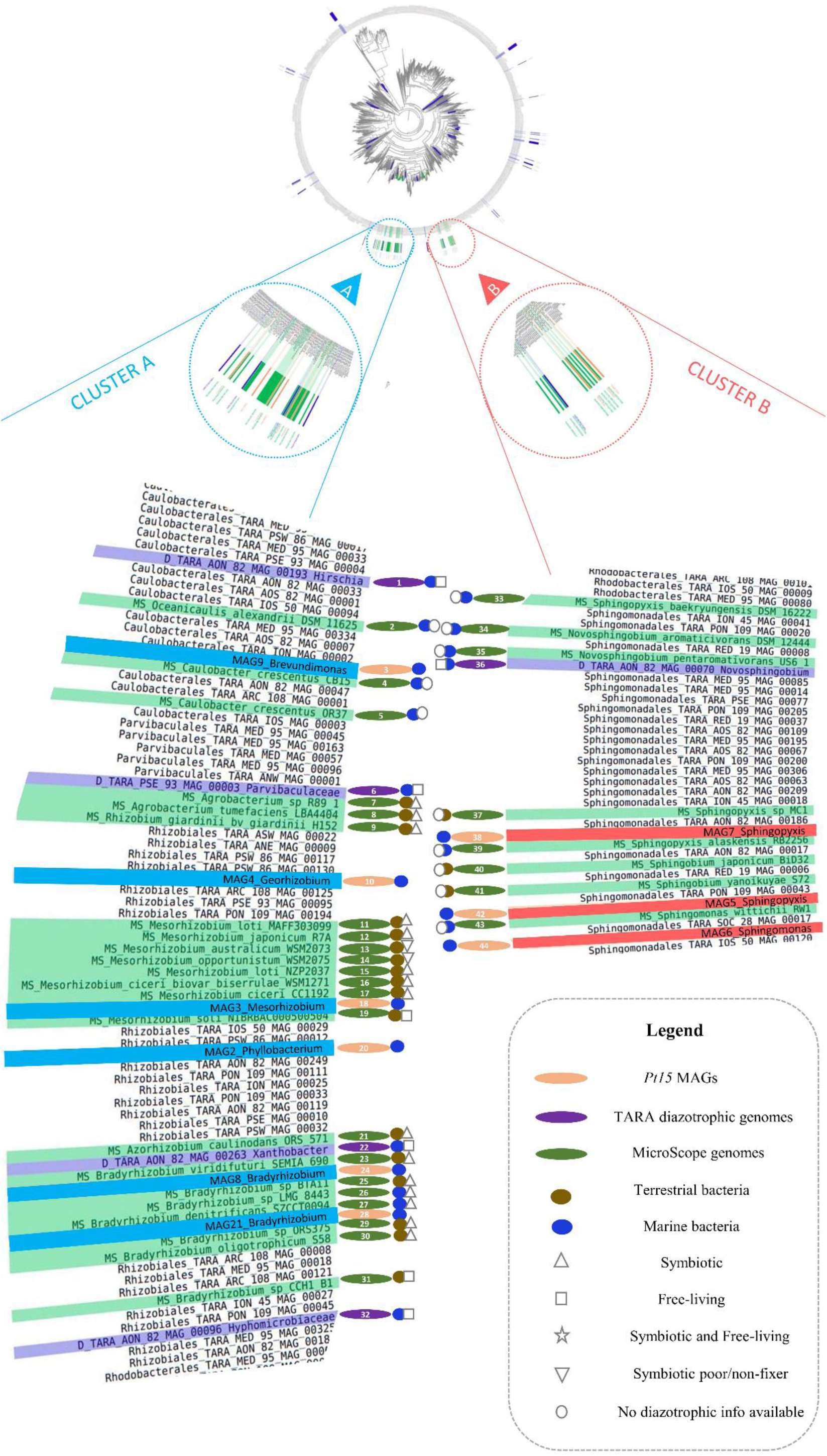
Phylogenetic tree of *Pt15* assembled bacterial genomes amongst 1888 TARA bacterial MAGs and MAGs from MicroScope. The phylogeny of 71 single copy core genes was built using Anvio platform. The upper circle represents a global view of the phylogeny built using a total of 1927 MAGs and genomes (1888 TARA MAGs, 30 terrestrial MAGs and 9 Pt15 MAGs). The bootstrap values represented from 0 – 1 with replicates of 100 and complete phylogeny can be accessed as an interactive at: https://ncdstree.univ-nantes.fr. The lower part of the figure represents a partial zoom-in on the two clusters where the Pt15 MAGs form groups in the tree with respect to other MAGs and genomes. Cluster A, left (blue highlighted text on the tree) consists of Pt15 associated MAGs: MAG9, MAG4, MAG3, MAG2, MAG8 and MAG21. Cluster B, right (red highlighted text on the tree) consists of Pt15 associated MAGs: MAG7, MAG5 and MAG6. Green highlighted text represents terrestrial bacteria, purple highlighted text represents diazotroph TARA MAGs (reported nifH gene in silico) and non-highlighted text/white background represents non-diazotroph TARA MAGs. Symbol legends, right lower-end denote information with bacterial genomes/MAGs on the tree. (Orange, oval): Pt15 MAGs, (purple, oval): TARA diazotrophic genomes (green oval): MicroScope genomes, (brown, circle): terrestrial bacteria, (blue, circle): marine bacteria, (triangle): symbiotic, (box): free-living, (star): symbiotic and free-living, (inverted triangle): symbiotic and poor- or non-fixer and (circle): no diazotrophic information available. The numbers indicated on the oval symbols correspond to genome information in Table S6.

### Functional gene annotation of *Pt15* associated NCDs in nitrate deplete versus replete conditions

To understand the biological functions of the microbial community, genes from -N and +N treatments were assigned to functional categories (**Fig. 4, Table S4**). There was a significant enrichment of genes related to nitrogen fixation and metabolism in -N compared to +N condition. Typically, in -N, we exclusively identified *nifH* and *nifD,* two key components of the nitrogenase enzyme complex. Likewise, *nifX* and *nifT* were only detected in -N suggesting their critical role in nitrogen fixation. Part of the *nif* operon, *NifX* was reported to be important in FeMo-co maturation and its transfer to *NifDK* proteins [33, 34]. Other genes related to nitrogen fixation metabolism emerged from the analysis, namely *HemN*, *Fpr*, *FixS*, *FixA*, *FixH*, *GroEL*, *NifS* and *FixJ* (**Fig. 4**), some of which are part of the NIF regulon.

**Fig. 4.**
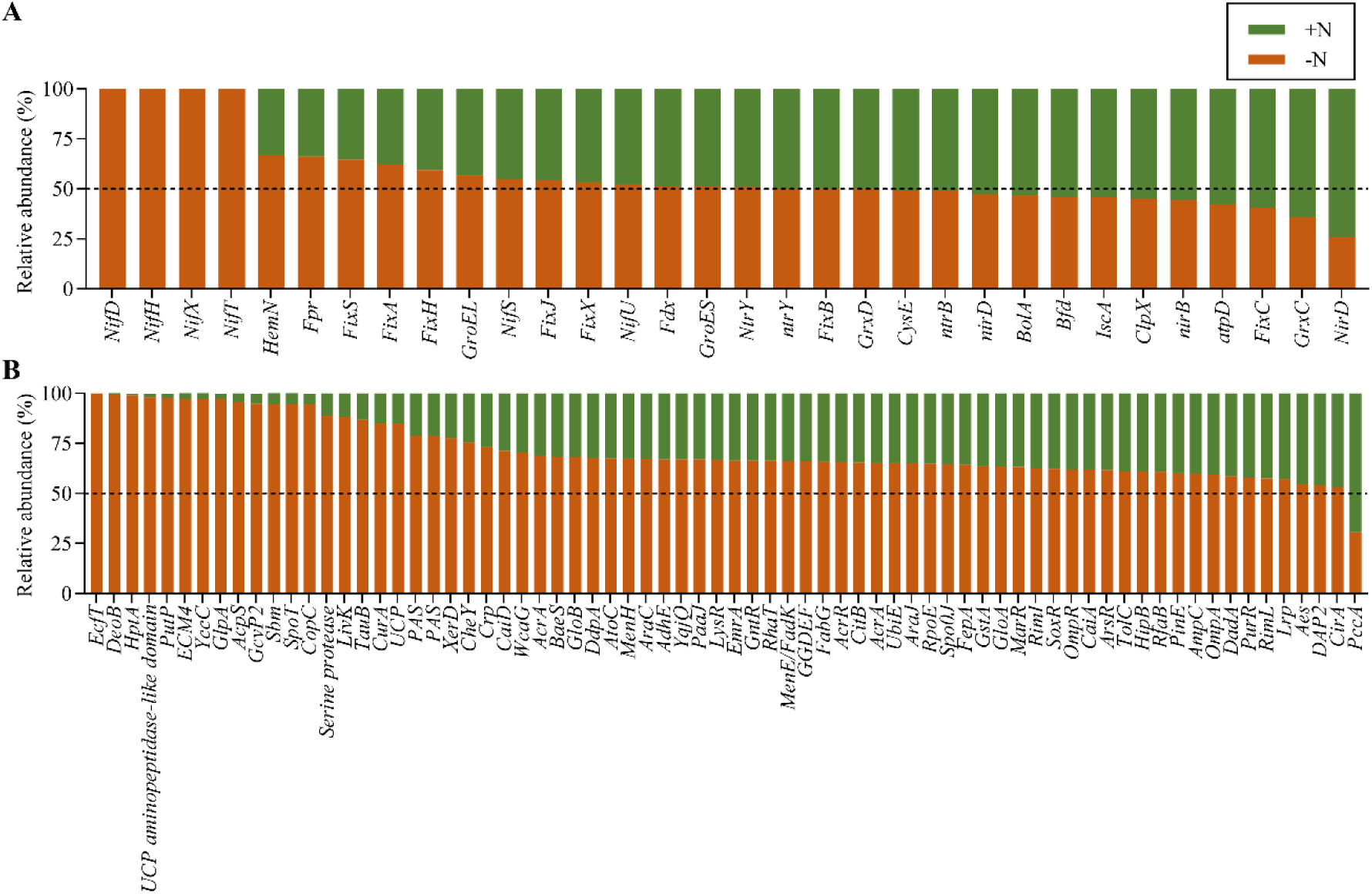
Functional gene annotation of *Pt15* associated NCDs in nitrate deplete versus replete conditions. A, Relative abundance of selected genes involved in nitrogen fixation and assimilation in nitrate deplete and nitrate replete conditions. B, illustration representing a combined view of 50 top abundant gene in nitrate deplete and replete condition with ordering representing top relatively abundant genes in nitrate deplete conditions. Dashed line indicates the relative percentage at 50%.

The abundance of several transporters was striking in -N, suggesting an important role in energy/substrate shuttling between the host and the bacterial community. Examples included an *EcfT* gene (energy-coupling factor transporter transmembrane) known for its function in providing the energy necessary to transport a number of different substrates [35], AcpS, an acyl carrier gene, which functions as a multi-carbon units carrier into different biosynthetic pathways [36], TauB, a taurine transporter that might play a role in providing taurine, which is produced by a large number of algae and known as a potential source of carbon, nitrogen and energy [37].

Stress responsive genes were clearly more represented in -N compared to +N including SpoT, which is a general bacterial stress response [38] that allows bacteria to increase its persistence in stressful conditions, here the lack of nitrogen. Other genes found almost exclusively in the starved bacterial community were the PAS domain containing genes (Per-ARNT-Sim) [39], which function as signal sensors by their capacity to bind small molecules, modulators and transducers. They are known to sense oxygen to prevent degradation of oxygen sensitive enzymes suggesting a crucial and protective role in -N condition where there is an enrichment in nitrogenase containing bacteria. Overall, functional analysis of MetaG data corroborated with NCDs enrichment in -N and nitrogen fixation.

### Geographical distribution and ecological prevalence of NCD-microalgae interactions

Next, we sought to assess the prevalence and importance of similar interactions in other microalgae providing further evidence that the association of NCDs with *P. tricornutum* is not an isolated case. First, we screened a collection of 14 xenic microalgae as described for *Pt15,* and identified five species of microalgae (35%) that grew in the absence of nitrogen in the growth medium suggesting the presence of diazotrophs (**Table S5**). Similar to *Pt15*, the retained microalgae were screened using different nitrogen deprived media under various conditions for diazotrophs isolation. *NifH* degenerate and 16S rRNA gene primers were used to identify the isolated bacterial colonies, which revealed five Alpha-proteobacteria including two species of *Thalassiospira sp*. (isolated from two different microalgae), *Stenotrophomonas maltophilia, Stappia* and *Rhodococcus qingshengii* suggesting that the occurrence of NCDs with microalgae may be relatively common. PCR detection using *nifH* degenerate primers was successful only in *S. maltophilia*. In all other bacteria, with the exception of *Stappia* and *S. maltophilia* which were not tested, a reduction of acetylene to ethylene was observed in the ARA assay **(Table S5)**. All isolated bacterial species grew in nitrogen free medium (**Table S5**). To provide additional support for the presence of nitrogen fixation in bacteria isolated from microalgae other than *P. tricornutum*, we carried out an ^15^N2 incorporation assay followed by LC-MS analysis on *Rhodococcus.* The results clearly indicated that the bacteria assimilated labeled nitrogen, confirming its ability to fix nitrogen (**Table S1**).

To maximize the number of species sequenced while maintaining cost-effectiveness, isolated NCDs were sequenced in pools of three bacteria using both Illumina and PacBio. Their genomes were successfully assembled, demonstrating high completeness (>87%, with two MAGs reaching 100%) and low contamination (**Table S3**). We then used MAGs isolated from *Pt15* and the other microalgae (**Table 1 and Table S3**) to reassess their phylogenetic relationship as described for *Pt15* isolated NCDs. The analysis revealed 3 more clusters (C, D and E) in close proximity to diazotrophic MAGs from the environment and/or MAGs belonging to diazotrophic containing orders [40] [41] [42] (**Figure S3**). An interactive figure is available at https://ncdstree2.univ-nantes.fr/.

Next, we assessed the occurrence of *Pt15* MAGs in the environment in surface water and Deep Chlorophyll Maximum layer (DCM) using *Tara* Ocean datasets [43, 44]. Given that *Pt15* MAGs had not been previously reported, the analysis to monitor the distribution for the surface and DCM waters was performed using the two phylogenetically closest TARA MAGs. The phylogeny was built in Anvio with 1888 TARA MAGs with inbuilt single copy genes HMM (Bacteria_71) and the five *Pt15* MAGs in which the *nifH* gene was identified. We used branch support and branch length as proxies to select the closely related TARA MAGS. The two TARA MAGs clustering with each individual *Pt15* MAG were selected (**Table S3**). With the exception of two *Pt15* MAGs, *Georhizobium* and *Sphingomonas*, which show weak branch support with one of the two TARA MAGs, all other MAGs are robustly supported as being closely related to their respective TARA MAGs in the phylogeny. Among the 130 sampling stations, 92 stations showed the occurrence of at least one of the TARA affiliated MAGs reflecting a high prevalence (**Fig. 5A, Figure S4**). *Pt15* clustering with *Tara* MAGs were more frequently occurring with 129 recorded stations. The distribution of these MAGs was extensive, spanning all latitudes from south Pacific, Indian and Atlantic Oceans to polar regions. The MAGs were found in both surface and DCM samples, mainly within the 0.8-5µm and 0.22-1.6/3µm size fractions (**Fig. 5A, Figure S4**). Interestingly, TARA MAGs affiliated to *Pt15* MAGs co-occurred in surface and DCM samples in most cases, although their dominance in surface samples was prominent (**Fig. 5B**).

**Fig. 5.**
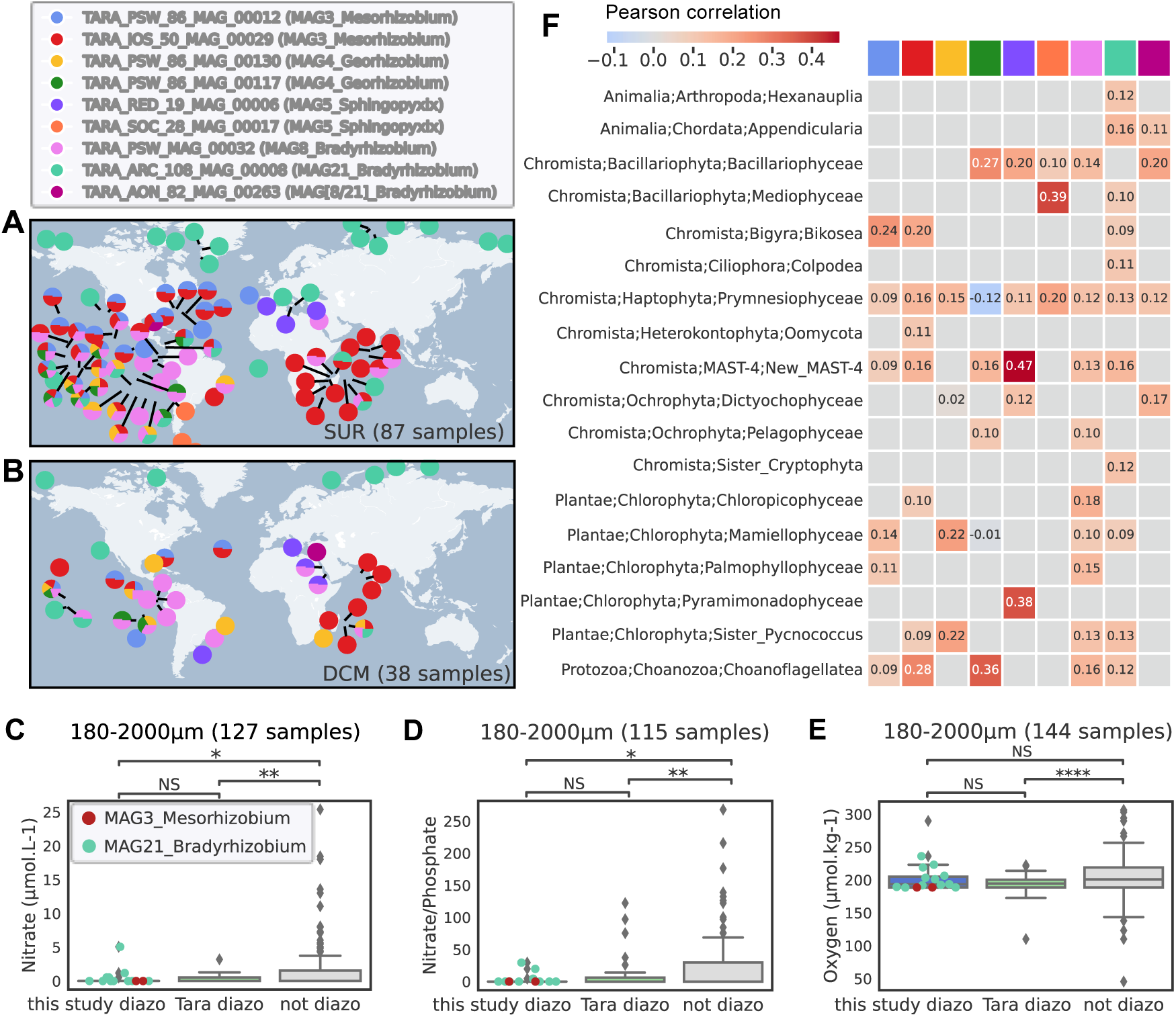
Geographical distribution and ecological relevance of isolated and assembled bacterial diazotrophs genomes correlating to TARA data. Geographical distribution of phylogenetically closest *Pt15* associated TARA Ocean MAGs in A, surface and B, DCM samples. C, D, relative abundance of the top 2 most occurring MAGs of *Pt15* bacterial community (C, D, *Mesorhizobium* MAG3, E, F *Bradyrhizobium* MAG8 respectively) in the environment in respect to nitrate, oxygen and iron concentrations. F, Co-occurrence of *Pt15* MAGs in the environment with different species of micro-eukaryotes inferred through FlashWeave, FDR <0.01)[100].

We then asked whether the occurrence of the MAGs isolated from *Pt15* correlated with environmental factors monitored during the *Tara* Oceans campaign and whether they co-occur with other organisms. We considered only two *Pt15* MAGs (*Mesorhizobium* MAG3 and *Bradyrhizobium* MAG21) to assess their correlation with three environmental factors used as indicators of diazotrophy, namely nitrate, nitrate phosphate ratio and oxygen concentrations. Iron concentrations from the ECCO2-DARWIN model [45] were considered, yet no distinct relationship was identified with the distribution of NCDs. Interestingly, TARA affiliated MAGs were detected in low nitrate and almost no detection above 5 µM (**Fig. 5C**). For both nitrate and N/P ratio, there are no differences in their occurrence with TARA diazotrophic MAGs, while these differences are significant with non-diazotrophic MAGs only in the larger size fraction of 180-2000 µm (**Fig. 5C and D**) suggesting an association with particles, aggregates or other organisms. This measure is in line with nitrate limitation as a marker of diazotrophs presence [46]. In a similar way, oxygen showed a consistent pattern when comparing this study’s MAGs with TARA NCDs, distinguishing these two categories from non-diazotrophic MAGs only in 0.22-1.6/3 µm, 0.8-5 µm and 180-2000 µm size fractions (**Fig. 5E and Figure S5**). By contrast to what is known about oxygen inhibition of nitrogenase, both MAGs seemed to thrive in relatively well oxygenated zones (200 µM) compared to less oxygenated areas or oxygen minimum zones (OMZ, 20-50 µM O2). This implies adaptive means by which these bacteria fix nitrogen such as creating micro-oxic environments through their interactions with other organisms.

Strikingly, a co-occurrence analysis revealed a clear association of the NCDs isolated from *Pt15* (**Fig 5H**) with micro-eukaryotes. Most of the *Pt15*-associated TARA MAGs were found to co-occur primarily with Haptophytes, Bacillariophyta, and Cryptophytes. The association with Bacillariophyta is consistent with their isolation from a diatom. While we did not demonstrate nitrogen fixation in these predicted associations, this analysis provides evidence for a probable nitrogen fixing interaction between *Pt15* MAGs and various species of primarily photosynthetic micro-eukaryotes.

## Discussion

Recent reports of NCDs dominance in the euphotic zones of the oceans questioned the long-held dogma that symbiotic N2 fixation is mainly carried by cyanobacteria, but also raised fundamental questions on how these presumably heterotroph NCDs can thrive in the sunlit ocean [12–14, 47]. The nitrogenase enzyme complex responsible for nitrogen fixation is sensitive to oxygen that irreversibly inactivates the enzyme. Therefore, diazotrophs evolved several mechanisms to protect nitrogenase from oxygen degradation including hyper-respiration, uncoupling of photosynthesis in space and time in photosynthetic diazotrophs, and attachment/interaction with microalgae or debris which were found to be hotspots of nitrogen fixation [17]. Using a multidisciplinary approach, the study described herein brings evidence of interactions for nitrogen fixation between microalgae, in particular the model diatom *Phaeodactylum tricornutum* and NCDs. After screening ten xenic accessions of *P. tricornutum* in the absence of a usable form of nitrogen, our study revealed that one accession, *Pt15*, interacts with nitrogen-fixing proteobacteria. Although low, this nitrogen fixation and uptake seemed to maintain the viability of the diatom and its associated bacteria. To compensate for the allocation of energy to the bacterial community, *Pt15* cells seemed to slow down growth. This diatom-NCDs interaction may reflect natural environments, where such interactions, in extreme conditions (low to no nitrogen) take place until favorable factors (e.g., nitrate repletion with upwelling or Sahara dust) permit the recovery and growth of cells, such as what we observed in our nitrate starved cultures when transferred to nitrate replete medium.

Metagenomic sequencing of the associated bacteria revealed not only one diazotrophic species but a group of nitrogen fixers predominantly Alpha-proteobacteria, mainly represented by Rhizobiaceae, Sphingomonadaceae and Xanthobacteraceae. All the isolated bacteria identified as diazotrophs grew in nitrogen free medium. The nitrogenase gene was detected in the majority of them, through either in silico analysis or by PCR amplification, further confirming the presence of diazotrophy in these bacteria. The *nifH* gene was found in most cases in the chromosome, except for *Georhizobium* where it was predicted to be plasmidic. ARA results indicated that nitrogen fixation rates in xenic *Pt15* under -N condition were lower than those observed in the positive controls and showed variability when compared to nitrogen fixation rates reported for NCDs in natural environments. These differences likely reflect the variability in environmental conditions across study sites. For example, in the Gotland Basin, NCD nitrogen fixation rates were estimated at approximately 7.6 nmol/L/day in surface waters and 0.44 nmol/L/day in deep waters [48]. In contrast, nitrogen fixation rates of NCDs in hypoxic waters of the Southern Californian Bight averaged 0.07 nmol/L/day, which is lower than those observed in the present study [49]. However, differences in monitoring techniques and experimental conditions, particularly between in situ measurements and laboratory cultures, must be considered when drawing comparisons [48–51]. The low nitrogen fixation rates observed in xenic *Pt15* and its associated bacterial community under nitrogen-depleted conditions are likely due to competition for fixed nitrogen by non-diazotrophic bacterial species. Another contributing factor may be the quiescent state of diatom cells induced by the absence of nitrogen in the culture. Our data indicate that diatom cells ceased division and likely maintained only minimal metabolic activity, which could have reduced the availability of photosynthates for associated bacteria. Quiescence is a well-documented survival strategy in certain microalgae under unfavorable environmental conditions, allowing them to save energy and sustain basic cellular functions until nutrient availability improves [52–54]. This was evident in *Pt15* cells, which resumed growth upon transfer to nitrogen-rich medium. If *Pt15* is not in an optimal physiological state to supply energy to its associated bacteria, nitrogen fixation rates will remain low, further exacerbated by the consumption of fixed nitrogen by the abundant non-diazotrophic bacteria present in the culture.

The nitrogen-fixing capability of the isolated bacteria was directly confirmed using ^15^N2 incorporation monitored by isotope analysis in LC-MS, leaving no doubt about the *Pt15* associated bacteria’s ability to fix nitrogen. This provides initial evidence that NCDs associated with the diatom are capable of fixing nitrogen, a process likely beneficial to other associated bacteria and potentially to the diatom, as long as it does not transition into a resting stage due to the extreme culture conditions, specifically the complete absence of bioavailable nitrogen. A recent study identified an NCD from the Rhizobiales order residing within a diatom of the *Halsea* genus, functioning as an endosymbiont [20]. The interactions we identified differ significantly from those with *Haslea* containing NCDs. The bacteria associated with *Haslea* have a much smaller genome, typically less than 1 Mb, whereas our NCDs have genomes about 7 Mb in size [20]. This discrepancy in genome size reflects the different types of interactions: in the obligate interaction, the bacteria have lost parts of their genome and depend on the host’s cellular machinery, whereas in the facultative interaction, the bacteria interact from a distance or within the phycosphere [55–58]. Our findings reflect the broad array of nitrogen fixation interactions, extending from obligatory symbiosis to epibiosis or remote interactions within the phycosphere, mirroring observations in nitrogen fixing interactions between Rhizobia and land plants. Bacteria associated with *Pt15*, identified as NCDs in this study, align with the distribution patterns of *Tara* NCDs in ocean regions characterized by low nitrate and nitrate phosphate ratio. Interestingly, their distribution and correlation with these environmental factors are observed only in larger size fractions. This observation is in accordance with findings from multiple studies that have reported similar diazotroph distributions under the aforementioned conditions. Indeed, diazotrophs have developed various strategies to counteract the irreversible degradation of the nitrogenase enzyme by oxygen [6, 7, 10, 59]. These strategies include forming associations with other organisms, among other mechanisms, as they seek out low-oxygen environments primarily facilitated by surrounding polymers.

Our study revealed that NCDs interaction with *Pt15* and other microalgae is likely not an isolated phenomenon and is more broadly distributed than previously anticipated. Our MAGs were detected in the environment clustering with previously identified terrestrial and marine NCDs, supporting diazotrophy and likely their ecological importance. These MAGs were identified at global scale in surface and DCM samples in size fractions enriched in microalgae, which supports the hypothesis of an interaction with them as evidenced by the co-occurrence analysis. This interaction might be taking advantage of hypo-oxic biomes microalgae and bacteria may provide with the secretion of extracellular polymers, mainly polysaccharides, that form biofilms ensuring low O2 diffusion [60–62]. This was suggested by our TEM analysis with the formation of pellicles around *Pt* cells and bacteria. Such pellicles were reported to form around two nitrogen fixing bacteria, *P. stutzeri* and *Azospirillum brasilense* in oxygenated surroundings implying the importance of oxygen stress in the formation of these structures [63]. The presence of these sac-like structures might have hindered diatom uptake of released ammonium leading to growth inhibition in the extreme condition tested in this study. Besides biofilm formation, this interaction may combine different strategies for both microalgae and NCDs to avoid/limit oxygen inactivation of nitrogenase such as a conformational switch that protects nitrogenase from O2 degradation [10, 64] or hyper-respiration [65] among other protective mechanisms that need further investigations. In line with our discovery of NCDs interaction with microalgae for nitrogen fixation, a recent study [66] suggested that anaerobic processes, including heterotrophic N2 fixation, can take place in anoxic microenvironments inside sinking particles even in fully oxygenated marine waters.

## Conclusions

Our screen under nitrogen-deprived conditions of different accessions of the model diatom *P. tricornutum* revealed several previously unidentified NCDs among which some were found to be related to terrestrial nitrogen-fixing bacteria, raising questions about their evolutionary origins. The NCDs associated with *P. tricornutum* enable the diatom to survive in nitrogen-deprived environments, indicating a relationship that supports its persistence. These NCDs remain in the phycosphere and do not penetrate diatom cells, suggesting that their interactions occur at a distance. Genome analyses of *P. tricornutum*-associated bacteria showed clustering with both terrestrial and marine NCDs, and these bacteria were found to be widely distributed across latitudes, co-occurring with photosynthetic microeukaryotes, including diatoms, haptophytes, and cryptophytes. The identification of different NCDs with a model diatom opens wide perspectives to investigate the role of individual and combined interactions of bacterial species with the host and within the bacterial community. Thus, *Pt15* accession with its associated NCDs represent a unique Eukaryotic-bacteria model system and an exciting opportunity to study molecular mechanisms underlying this interaction for nitrogen fixation and beyond.

## Methods

### Microalgae species used and culture media

#### Screening of Phaeodactylum tricornutum accessions

Eleven xenic *P. tricornutum* accessions were screened in presence (545 µM) and absence of nitrate in Enhanced Artificial Sea Water (EASW) in order to select xenic *Pt* ecotype growing in the absence of nitrate. The starting concentration for all accessions, measured under the microscope using Malassez counting chamber, was 100,000 cells per ml. The cells were counted and washed twice with nitrate-free EASW in order to remove all residual nitrate and resuspended in respective media for screening over a period of one month. A growth curve was made by counting the cells in periodic time intervals. The cultures were grown at 19°C with a 12/12 hours photoperiod at 30 to 50 µE/s/m2. A cocktail of antibiotics (Ampicillin 100 µg/µl, Chloramphenicol (14 µg/µl) and Streptomycin 100 µg/µl) was used to make *Pt15* axenic in liquid EASW culture medium with refreshment of antibiotic containing medium every 5 days. *Pt15* cells were regularly monitored for axenicity using peptone-rich medium and microscopy. Additionally, axenicity was further validated by culturing the putatively axenic *Pt15* in -N medium. If the *Pt15* cells died under these conditions, it would confirm the absence of associated bacteria.

#### Screening of other microalgae

A total of fourteen microalgae were screened in nitrate-deplete and nitrate-replete enhanced artificial sea water for growth (Table S5). About 100,000 cells were counted, resuspended in their respective media and grown as described above. Their growth was assessed for a period of two weeks at regular intervals.

### Acetylene reduction assay

Acetylene reduction assay (ARA) [21] was used to quantify the conversion of acetylene to ethylene (C2H4) which allows the assessment of nitrogenase activity. ARA was conducted on the following samples: two positive controls, *Azotobacter. vinelandii* (NCIMB 12096), *Pseudomonas stutzeri* BAL361, xenic *Pt15* grown in NFM [67, 68] or nitrogen replete medium, all incubated at three different temperatures (19, 25, 30°C). Axenic *Pt15* culture grown in NFM medium + 5 sugars (final concentration of 1% each: D-glucose, D-sucrose, D-fructose, D-galactose and D-mannose) was used in every measurement as a negative control to ensure there was no endogenous ethylene produced. All the cultures were grown at their respective growth media and conditions. After reaching an appropriate O.D. between 0.2-0.3, 1 ml of the bacterial cells were collected in an Eppendorf’s tube and centrifuged at11,000 rpm for 10 minutes at RT to remove the medium. The cells were re-suspended in NFM with five sugars and grown in their respective medium. Sugar was not used for culturing xenic *Pt15*. When the bacterial growth reached an O.D. of 0.2-0.3, the cultures were prepared for acetylene reduction assay. The cultures were secured with a rubber stopper in order to make them air tight. Ten percent of the air was removed from the culture flasks and supplemented with 10% acetylene gas. The cultures were incubated at their appropriate temperatures for a period of 72 hours before monitoring ARA with gas chromatography. A volume of 500 µl was taken from the headspace of the samples using a Hamilton gas tight syringe (SYR 500 ul 750 RN no NDL, NDL large RN 6/pk) and injected into an Agilent gas chromatograph model 7820 model equipped with a Flame Ionization Detector (GC-FID) and using Hydrogen as carrier gas at a constant flow of 7.7 mL/min. Samples were injected into a split/splitless injector, set at 160 °C in splitless mode. A GS-Alumina capillary column, 50 m × 0.53 mm × 0.25 μm (Agilent, 115-3552) was used. The programmed oven temperature was at 160 C° (isotherm program) during 2 min. The FID detector was set to 160 C° . A standard curve was made with ethylene gas using 5 volumes (100, 150, 200 µl). The raw data generated from the GC provides value of y, Area [pA*s]. This value is supplemented in the slope-intercept formula, y = mx + c, with ‘m’ and ‘c’ representing slope and constant values respectively, denoted in the calibration curve. The amounts of ethylene were calculated as described previously [69]. Acetylene reduction rates were converted to fixed N using the common 3:1 (C2H2 :N2) molar conversion ratio [70, 71].

### Particulate organic nitrogen measurements

One liter of 200,000 cells/ml culture of xenic *Pt15* was grown in nitrate-free EASW for over a period of 80-days at 19°C and 12h/12h light/dark photo period at 70µE/s/m2. Six time points were used for harvesting samples: 1, 4, 12, 59, 76 and 80 day(s). The cells were counted at each time point using a cell counter. Twenty ml of *Pt15* culture were harvested and injected through a sterile (treated at 450 deg/4hrs) Whatman GF/F glass microfibre filter paper to collect *Pt15* cells and the filters were dried overnight in 60°C incubator and stored at -80°C. The 20 ml filtrate from the cultures was passed through 0.22 µm filters (Sigma ca. GVWP10050), in order to remove bacterial cells and stored at -80°C, before being processed for particulate organic nitrogen measurement analysis. EASW with 545 µM nitrate, 0 µM nitrate and three heat-treated Whatman GF/F glass microfibre filters were used as blanks for comparison with test samples.

### Test for incorporation of the ^15^N isotope

Stationary cultures (30 mL) of either the diatoms, *Phaeodactylum tricornutum Pt15* Xenic Plus nitrate (545 µM)*, P. tricornutum Pt15* Xenic Minus nitrate (0 µM)*, P. tricornutum Pt15* Axenic Plus nitrate (545 µM) or bacteria, *Bradyrhizobium MAG21*, *Methylobacterium*, *Azotobacter vinalendii*, *Pseudomonas stutzeri BAL361*, *Escherichia coli DH5a* and *Rhodococcus* were put under a ^15^N2-(98 atom %, Sigma-Aldrich, Steinheim, Germany) atmosphere (20 mL). For this, a 50 mL syringe (B Braun Omnifix®) without needle was used to take up 30 mL of the culture, followed by 20 mL of 15N2 from a lecture bottle. The syringes were immediately sealed with a plastic valve and parafilm to prevent leakage of the gas. Diatom cultures were incubated at 18 °C under a 14:10 h light: dark regime with an illumination of 60-65 µmol photons m^-2^ s^-1^. Bacterial cultures were incubated at 100 rpm and 30 °C, except for *E. coli DH5a* which was incubated at 37 °C. After 2.5 days the diatom cultures were filtered (1.2 µm pore size, GF/C, Whatman, GE Healthcare, Little Chalfont, United Kingdom). The cells were washed off the filter using 5 mL ice-cold methanol/ethanol/chloroform 1:3:1 (methanol: HiPerSolv Chromanorm, VWR Chemicals, Dresden, Germany; ethanol: HiPerSolv Chromanorm, VWR Chemicals, Dresden, Germany; chloroform: SupraSolv, VWR Chemicals, Dresden, Germany) and the resulting extracts were treated in an ultrasonic bath for 10 min and centrifuged (16.100 x g, 20 min, 4 °C). The supernatant was stored at −20 °C until analysis. Bacterial cultures were centrifuged (3.893 x g, 20 min, 4 °C) and the cells were resuspended in 5 mL of methanol/dichlormethane/ethylacetate 10:2:3 (dichlormethane: SupraSolv, VWR Chemicals, Dresden, Germany; ethylacetate, HiPerSolv Chromanorm, VWR Chemicals, Dresden, Germany). The suspensions were sonicated (5 min, 9 cycles, 80-85 %) and afterwards centrifuged (3893 x g, 20 min, 4 °C). The supernatant was stored at −20 °C until further use.

### Test for uptake of ^15^N-L-valine-by *P. tricornutum*

To stationary cultures (40 mL) of either *Phaeodactylum tricornutum Pt15* Xenic Plus nitrate (545 µM) or *P. tricornutum Pt15* Xenic Minus nitrate (0 µM), 100 µL of an aqueous ^15^N-L-valine-solution (98 atom % ^15^N, Merck, Saint Louis, USA, 50 µmol L^-1^ in medium) were added to reach a final concentration of 125 nm L^-1^. The cultures were then incubated, extracted and the samples stored as described in “*Test for incorporation of the ^15^N isotope”*.

### Analysis of endometabolomes with LC-MS

Cell extracts were dried in a nitrogen stream at room temperature and resuspended in either 200 µL methanol for diatom extracts or methanol/acetonitrile/water 5:9:1 (acetonitrile: HiPerSolv Chromanorm, VWR Chemicals, Dresden, Germany; water: CHEMSOLUTE, TH.GEYER, Renningen, Germany) for bacterial extracts. Metabolites were separated on a Dionex Ultimate 3000 UHPLC coupled to a Q-Exactive Plus Orbitrap mass spectrometer (Thermo Scientific, Bremen, Germany). Chromatographic separation was performed on a SeQuant ZIC-HILIC column (5 µm, 200 Å, 150 × 2.1 mm, Merck, Germany) equipped with a SeQuant ZIC-HILIC guard column (20 × 2.1 mm, Merck, Germany) at 25 °C using water with 2 % acetonitrile and 0.1 % formic acid as eluent A and 90 % acetonitrile with 10 % water and 1 mmol L^-1^ ammonium acetate as eluent B. Full scan measurements (*m*/*z* 80-1200, resolution 70k, automatic gain control target 3E^6^, maximum ion injection time 200 ms) were performed in positive and negative ion mode using a heated electrospray ionization source to generate molecular ions. The method duration was 14.5 min with an MS runtime from 0.5 min to 9.0 min, a flow rate of 0.6 mL min^-1^ and a gradient as follows: 85 % B (0.0-4.0 min), linear gradient to 100 % A (4.0-5.0 min), 100 % A (5.0-9.0 min), linear gradient to 100 % B (9.0-10.0 min), 100 % B (10.0-12.0 min), linear gradient to 85 % B (12.0-12.5 min), 85 % B (12.5-14.5 min). Amino acid standards were run separately for comparison of retention time and mass spectra. Resulting raw data was analyzed with FreeStyle (Thermo Fisher Scientific).

### Microscopy

#### Light microscopy

Living cells grown on agar plates were used directly for the preparations. Cells were taken directly from agar plates without prior treatment, transferred onto glass slides, and covered with a cover slip before observation. Samples were observed with differential interference contrast (DIC) microscopy by BX53 microscope (Olympus, Japan) and Axiocam 705 color (Carl Zeiss, Germany).

#### Transmission electron microscopy

Samples were prepared as described previously [72]. Embedded samples were cut for making pale yellow ultra-thin sections (70-80nm thickness) with a microtome EM UC6 (Leica Microsystems, Germany). Sections were stained with TI blue (Nisshin EM, Japan) and lead citrate for inspection with an electron microscope JEM-1011KM II (JEOL, Japan).

### Bacterial isolation and culture media used

A total of eight media recipes were used for bacteria isolation and culture under different conditions, all summarized in Supplementary File 1. All isolated bacteria were checked for purity by repeated streaking and 16S sequencing. Bacterial species were maintained in 20% glycerol at - 80°C for long-term storage.

### DNA extraction and PCR amplification of marker genes

For initial identification of bacteria, the isolated colonies were picked under sterile conditions using filter tips, resuspended and mixed thoroughly in 50 µl nuclease-free water (NFW). The suspension was heat/cold treated at 95°C following 4°C, 10 min each and twice for cell lysis. Post-treatment, an additional 50 µl NFW was added to the suspension for a total volume of 100 µl. Five µl of the colony DNA was used as template for PCR.

Wizard® Promega Genomic DNA extraction kit was used for extracting high-quality genomic DNA from bacterial cells. The bacteria were grown in their appropriate media until the culture optical density (OD) reached 0.3. The cells were harvested and washed in 1X PBS twice. The cell pellets were resuspended in 480 µl 50 mM EDTA and treated with 120 µl of 20 mg/ml lyzozyme (VWR cat no. 0663-5G), incubated at 37°C for 60 min in order to lyze possible gram-positive bacteria. Post incubation, the cells were centrifuged and the supernatant was removed. Nuclear Lysis solution, 600 µl (Promega cat no. A7941), was used to re-suspend and gently mix the pellet for incubation at 80°C for 5 min. The suspension was allowed to cool at room temperature (RT) for 10 min before adding 1 µl RNAse at 10 mg/ml (VWR cat no. 0675-250MG) and incubation at 37 for 60 min. After incubation, the suspension was cooled at RT for 10 min. Protein lysis solution, 200 µl (cat no. VWR 0663-5G) was added, vortexed and incubated on ice (4°C for 5 min). The mixture was centrifuged and the clear supernatant was transferred to a clean Eppendorf with 600 µl RT isopropanol and the tube was mixed gently before being centrifuged. The cell pellet was washed with freshly prepared 70% ethanol and re-centrifuged. The ethanol was carefully removed without disturbing the pellet and air dried before resuspension in appropriate amount of NFW. The quality of the DNA was checked using Nanodrop 260/280 and 230/260 ratios and on 1% agarose gel. The identity of the bacteria was re-confirmed using 16S rRNA gene amplification and Sanger sequencing.

For the *nifH* PCR amplification, the reaction mixture included 2.5 μL of DreamTaq Green Buffer (1X), 0.1 μL of BSA (0.08 µg/µL), 2 μL of each primer (10 μM), 0.2 µL of DreamTaq Green Polymerase (5 U/µL), 0.5 µL of dNTP mix (10 mM), 3 µL of DNA template (normalized to 10 ng/µL), and H2O adjusted to a total volume of 25 μL. The thermocycling protocol started with an initial denaturation step at 95°C for 5 minutes, followed by 40 cycles of denaturation at 95°C for 30 seconds, annealing at 53°C for 30 seconds, and extension at 72°C for 30 seconds, and a final extension step at 72°C for 7 minutes. Following the reaction, amplification results were assessed through 1% agarose gel electrophoresis, and DNA bands of interest were isolated using the NucleoSpin® Gel & PCR Cleanup Kit from Macherey-Nagel. The extracted DNA underwent Sanger sequencing at Eurofins Scientific. Sequences were analyzed using Geneious. For species identification, 16S PCR fd1-rd1, variable regions PCR V1V2 V3V4 and V5V7 were used. The reaction was performed with 35 cycles of 95 °C (1 min), 55 °C (1 min) and 72 °C (1 min). All primer sequence details are reported in supplementary Table 7.

### Sample preparation and DNA extraction for Illumina and PacBio sequencing

Duplicate with a starting concentration of 200,000 cells/ml of 3-liter culture volumes of xenic *Pt15* in nitrate-replete and nitrate-deplete enhanced artificial sea water were grown for a period of 2 months (60-days) at 19°C and 12h/12h light/dark photoperiod at 70µE/s/m2 before collection for sequencing. The xenic *Pt15* cells were filtered through a 3 µm filter (MF-millipore hydrophobic nitrocellulose, diameter 47 mm (cat no. SSWP04700) to retain the cells and allow passage of the filtrate containing the bacterial community. The filtrate was filtered twice in order to avoid contamination of the bacterial cells with residual *Pt15* cells. The double filtered bacterial community was collected in 0.22-micron filter (MF-millipore hydrophobic nitrocellulose, diameter 47 mm (cat no. GSWP04700) and resuspended in 1X PBS. DNA was extracted for metagenome sequencing using Quick-DNA fungal/bacterial mini prep kit (cat no. DG005) and according to manufacturer instructions. The bacterial DNA quality was checked using 1% agarose gel to assess intact DNA and quantified using Qubit broad range DNA kit, before being sequenced. Additionally, 16S rRNA sequencing was performed to assess the presence of bacterial DNA and 18S rRNA PCR was run as a negative control to inspect absence of *P. tricornutum* DNA. When bacterial isolates were available, they were sequenced either as a single species or in a pool of 3 bacteria using Illumina and PacBio.

### Data processing

#### Metagenomic analysis

Two different strategies were used to analyze the metagenomic reads: (i) a “Single assemblies” strategy, where assembly and binning was performed for each sample individually. This strategy is known to produce less fragmented assemblies with a limitation of chimeric contigs formation [73]. (ii) a co-assembly strategy was also performed to complete the results with additional likely lower abundant MAGs. The idea is to pool all the available samples reads and build a unique and common assembly for all samples. Six samples from 2 experiments were used: one with sequences obtained at 2g/L salinity (in +N and -N conditions, no replicate) and the other one at 20 g/L salinity with duplicates for each of the two conditions. Both datasets are similar.

##### Single assembly strategy

DNA raw reads were analyzed using a dedicated metagenomic pipeline ATLAS v2.7[73]. This tool includes four major steps of shotgun analysis: quality control of the raw reads, assembly, binning, taxonomic and functional annotations. Briefly, quality control of raw sequences is performed using utilities in the BBTools suite (https://sourceforge.net/projects/bbmap/): BBDuk for trimming and removing adapters, BBSplit for decontamination. In our case *Phaeodactylum tricornutum* reference genome (accession GCA_000150955.2), including mitochondria and chloroplast sequences, was used for this specific step. Quality-control (QC) reads assemblies were performed using metaSPADES[74] with ATLAS default parameters. The assembled contigs shorter than 500 bases length and without mapped reads were removed to keep high-quality contigs.

Metagenome-assembled genomes (MAGs) were obtained using two distinct binners : maxbin2[75] and metabat2[76]. The bins obtained by the different binning tools were combined using DASTool[77] and dereplicated with dRep[78] using default parameters. Completeness and contamination were computed using CheckM[79]. Only bins with a completeness above 50 % and a contamination rate under 10 % were kept. Prediction of open reading frames (ORFs) was performed, at the assembly level (not only at the MAGs level) using Prodigal[80]. Linclust[81] was used to cluster the translated gene product obtained to generate gene and protein catalogs common to all samples.

##### Co-assembly strategy

QC reads and co-assembly were analyzed using a shotgun dedicated sequential pipeline MetaWRAP v1.3.2[77]. Briefly, QC reads are pulled together and addressed to the chosen assembler metaSPAdes[74] using MetaWRAP default parameters. Metagenome-assembled genomes (MAGs) were obtained using two distinct binners: maxbin2 and metabat2 and the bins obtained were combined using an internal MetaWRAP dedicated tool. Completeness and contamination were computed using CheckM[79]. Only bins with a completeness above 50 % and a contamination rate under 10 % were kept. MetaWRAP also includes a re-assemble bins module that collect reads belonging to each bin, and then reassemble them independently with a “permissive” and a “strict” algorithm. Only improved bins are kept in the final set. Using this strategy, we were able to add an additional MAG.

### MAGs relative abundances

The QC reads were mapped back to the contigs, and bam files were generated to organize downstream analysis. Median coverage of the genomes was computed to evaluate MAGs relative abundance estimations.

### Taxonomic assignation of metagenomic data

We utilized anvio version 7.2[29] for taxonomic profiling of the *P. tricornutum* associated metagenomes in nitrate deplete and replete condition. The workflow uses The Genome Taxonomy Database (GTDB)[23] to determine the taxanomy based on 22 single copy core genes (scgs, anvio/anvio/data/misc/SCG_TAXONOMY). Sequence alignment search was done using DIAMOND [82].

### Hybrid assembly of MAGs

#### Assembly

Metagenomic assembly of Illumina technology reads does not yield complete genomes due to the difficulty of assembling repetitive regions [83]. Whenever Illumina and PacBio reads were available, we used a strategy to improve MAGs assembly by combining both type of reads. Metagenomic assembly of Illumina paired-end and PacBio reads post quality trimming were assembled using metaSPAdes [74] or SPAdes(v3.15.5) with option flag (--meta). To validate the newly assembled genomes, we verified that the long reads covered regions that were not supported by the mapped Illumina sequences only and that we get high quality bins. We obtained more contiguous genomes in which some regions had not been supported by the assembly of Illumina reads only.

#### Binning

The contigs were grouped and assigned to the individual genome. To do this, we used MaxBin2 and MetaBAT2 which are clustering methods based on compositional features or on alignment (similarity), or both[84].

#### Refinement

Bins from MaxBin2 and MetaBAT2 were consolidated into one more solid set of bins. To do this we used Binning_refiner which improves genomic bins by combining different binning programs available at: https://github.com/songweizhi/Binning_refiner [85].

#### Taxonomic Assignation

GTDB-Tk has already been independently and positively evaluated for the classification of MAGs[86]. After consolidating the bins in the refinement step, we determined the taxonomy of each bin. The gtdbtk classify_wf module was used in this work using a GTDB database https://github.com/ecogenomics/gtdbtk [87].

### Gene annotation of metagenomic data and MAGs

Gene annotation of metagenomics data in two conditions was done using anvi’o version 7.2. Three databases were used for a robust gene observations: COG20[88], KEGG[89] and Prokka (https://github.com/tseemann/prokka). DIAMOND was used to search NCBI’s Database. In built, anvi’o program annotates a contig database with HMM hits from KOfam, a database of KEGG Orthologs (KOs, https://www.genome.jp/kegg/ko.html). For Prokka, the annotation was done externally and the gene calls were imported to anvi’o platform for the individual metagenome contigs databases.

### Phylogenomic analysis of MAGs

The phylogenomic tree was built using anvi’o -7.1 with database of 1888 TARA ocean MAGs (contig-level FASTA files) submitted to Genoscope (https://www.genoscope.cns.fr/tara/). Additional closest terrestrial and marine genomes (Supplementary Table 6) to the individual MAGs were selected using MicroScope database (https://mage.genoscope.cns.fr/microscope/) using genome clustering, utilizing MASH [90] for ANI (Average nucleotide identity) for calculating pairwise genomic distancing, neighbor-joining (https://www.npmjs.com/package/neighbor-joining) for tree-construction and computing the clustering using Louvain Community Detection (https://github.com/taynaud/python-louvain). anvi’o utilizes ‘Prodigal’[80] to identify open reading frames for the contig database. The HMM profiling for 71 single core-copy bacterial genes (SCGs) is done using in-built anvi’o database, Bacteria_71 (https://doi.org/10.1093/bioinformatics/btz188) and utilizes MUSCLE[91] for sequence multiple alignment. The bootstrap values are represented from 0 – 1 with replicates of 100.

### Plasmid prediction and annotation

Plasmids were predicted from MetaG data using the following plasmid assembly tools: SCAPP[92], metaplasmidSPAdes[93] and PlaScope[94]. Predicted plasmids were annotated using Dfast[95], eggNOG-mapper v2[96] and Prokka 1.14.6[97].

### *Tara* Oceans MAGs biogeography and environmental preferences

Previously reconstructed MAGs from Tara Oceans metagenomes and respective abundance count tables and fasta files were downloaded from open-source released *Tara* Oceans databases at https://www.genoscope.cns.fr/tara/[13, 98]. Data included 2601 MAGs of which 1888 prokaryotes and 713 eukaryotes MAGs in 937 samples at 2 depths (Surface and Deep Chlorophyll Maximum) and 5 size fractions (0.22-3μm, 3-20μm, 0.8-5μm, 20-200μm, 0.8-2000μm, 180-2000μm) [99]. Data included 87 and 38 samples respectively at surface and DCM depths and 6 size fractions (0.22-3μm, 3-20μm, 0.8-5μm, 20-200μm, 180-2000μm).

The biogeographical occurrences of TARA MAGs phylogenetically close to Pt15 were projected onto global maps for both surface and DCM depths, either by grouping the 5 size fractions (Figure 5), or separately by size fraction (fig. S4).

Tara Oceans physico-chemical parameters were downloaded from Pangaea (*38*). MAGs TSS relative abundances were projected and analyzed in relationship with ammonium (µmol/L), chlorophyll A (mg/m3), iron (µmol/L), nitrite (µmol/L), nitrate (µmol/L), phosphate (µmol/L), nitrate/phosphate, oxygen (µmol/kg), salinity (PSS-78), silicate (µmol/L), temperature (°C) and photosynthetically available radiation (mol quanta/m2/day).

### Co-occurrence network inference

Co-occurrence network inference was performed using FlashWeave v0.18.0 [100] with Julia v1.5.3, for eukaryotic and prokaryotic Tara MAGs linked to *Pt15* MAGs. To account for the compositional nature of MAGs abundance data, a multiplicative replacement method [101] was applied to replace zeros, and a Centered Log Ratio transformation [102] was applied separately on both prokaryotic and eukaryotic abundance matrices. FlashWeave parameters used were max_k=1, n_obs_min=10, heterogeneous=False and normalize=False.

## Availability of data and materials

All data needed to evaluate the conclusions in the paper are present in the paper and/or the Supplementary Materials. All sequencing data and generated assemblies will be available under the BioProject PRJNA923279.

## Acknowledgements

We acknowledge Clara Guillouche and Irene Romero Rodriguez for their technical assistance with PCR screening and Romain le Balch for his help with GC runs. LT acknowledges support from the region of Pays de la Loire (ConnecTalent EPIALG project), Epicycle ANR project (ANR-19-CE20-0028-02) and µAlgaNIF France-Japan International Research Project. UC was supported by grant 998UMR6286 Connect Talent EpiAlg from Région Pays de la Loire to LT. SC acknowledges support from the CNRS MITI through the interdisciplinary program *Modélisation du Vivant* (GOBITMAP grant), and the H2020 European Commission project AtlantECO (award number 862923). We are grateful to the bioinformatics core facility of Nantes University (BiRD Biogenouest) for its technical support. The LABGeM (CEA/Genoscope &CNRS UMR8030), the France Génomique and French Bioinformatics Institute national infrastructures (funded as part of Investissement d’Avenir program managed by Agence Nationale pour la recherche, contrats ANR-10-INBS-09 and ANR-11-INBS-0013) are acknowledged for support within the Microscope annotation platform.

## Contributions

LT conceived and designed the study. UC performed most of the experiments. SM contributed to the ARA assay and the screening. LJLA assisted with the screening and performed bacterial isolation. TC, CT, EM, UC performed bioinformatics analysis. MG performed bioinformatics analysis and generated figure 5. TL conducted the PON assay. AT performed the EM study. IL assisted with the ARA assay. EJF performed 15N assay. GP supervised EJF and provided guidance on the 15N assay. SC supervised MG. UC, TC, CT, MG, SC, EJF, GP, AT, TL and LT analysed and interpreted the data. LT wrote the manuscript with methodological input from UC, MG, EF, AT, TL, EM and TC. LT supervised and coordinated the study. All authors read, edited and approved the manuscript.

## Competing interests

The author(s) declare no competing interest.

## Supplementary figures and tables legend

**Figure S1.**
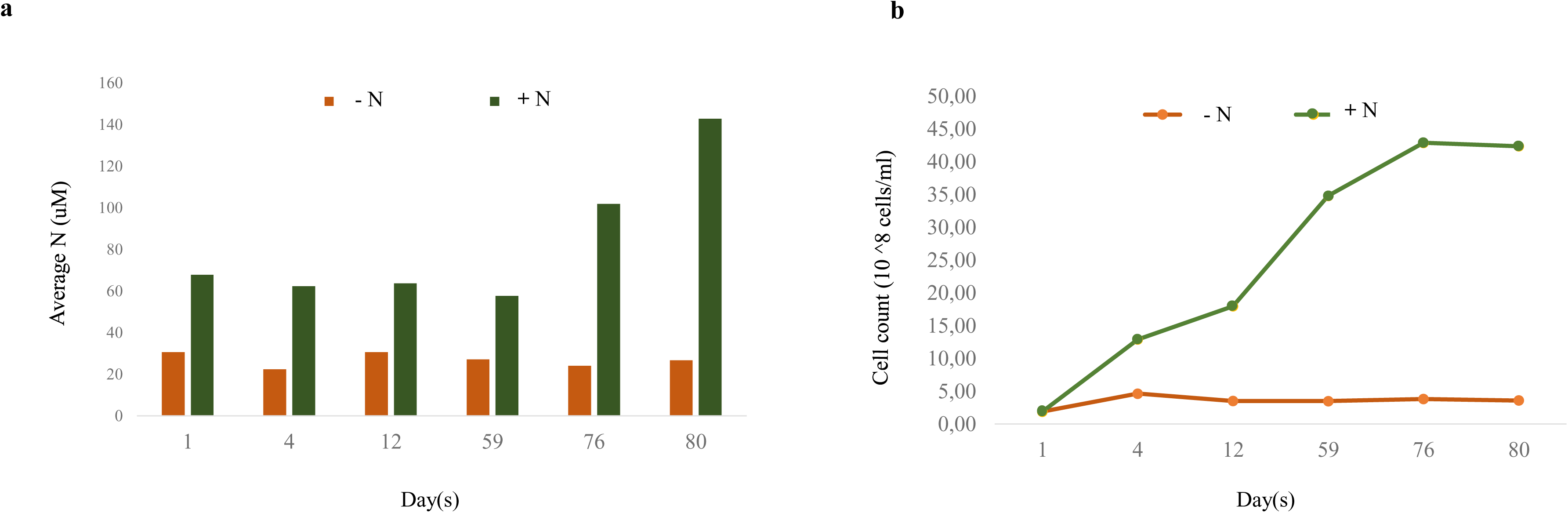
**Particulate Organic Nitrogen (PON) measurements**. a, PON kinetic and b, corresponding xenic *Pt15* biomass for 6 time points over a period of 80 days in nitrate deplete and replete conditions.

**Figure S2.**
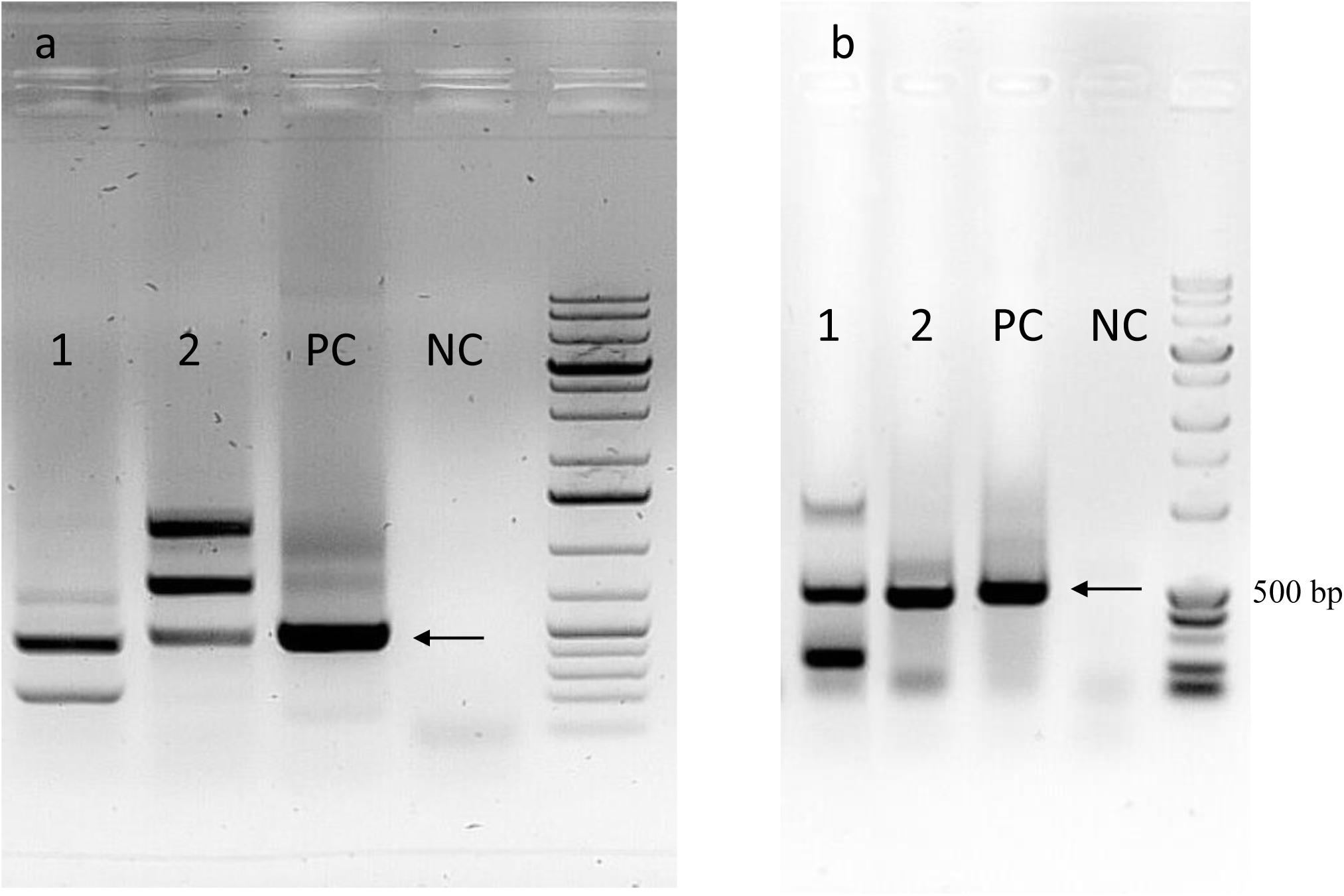
Agarose gel pictures of PCR amplified products. A, amplification of *nifH* gene in isolated bacteria using degenerate primers. 1, *Stenotrophomonas maltophilia* MAG20, 2, *Bradyrhizobium* sp. (MAG 21), 3, Positive control (PC) *Azotobacter vinelandii,* Negative control (NC). B, 1, *Paracoccus sp*., 2, *Bradyrhizobium sp.,* (MAG 21), *Azotobacter vinelandii* (PC), Negatice control (NC). A black arrow indicates the *nifH* PCR band.

**Figure S3.**
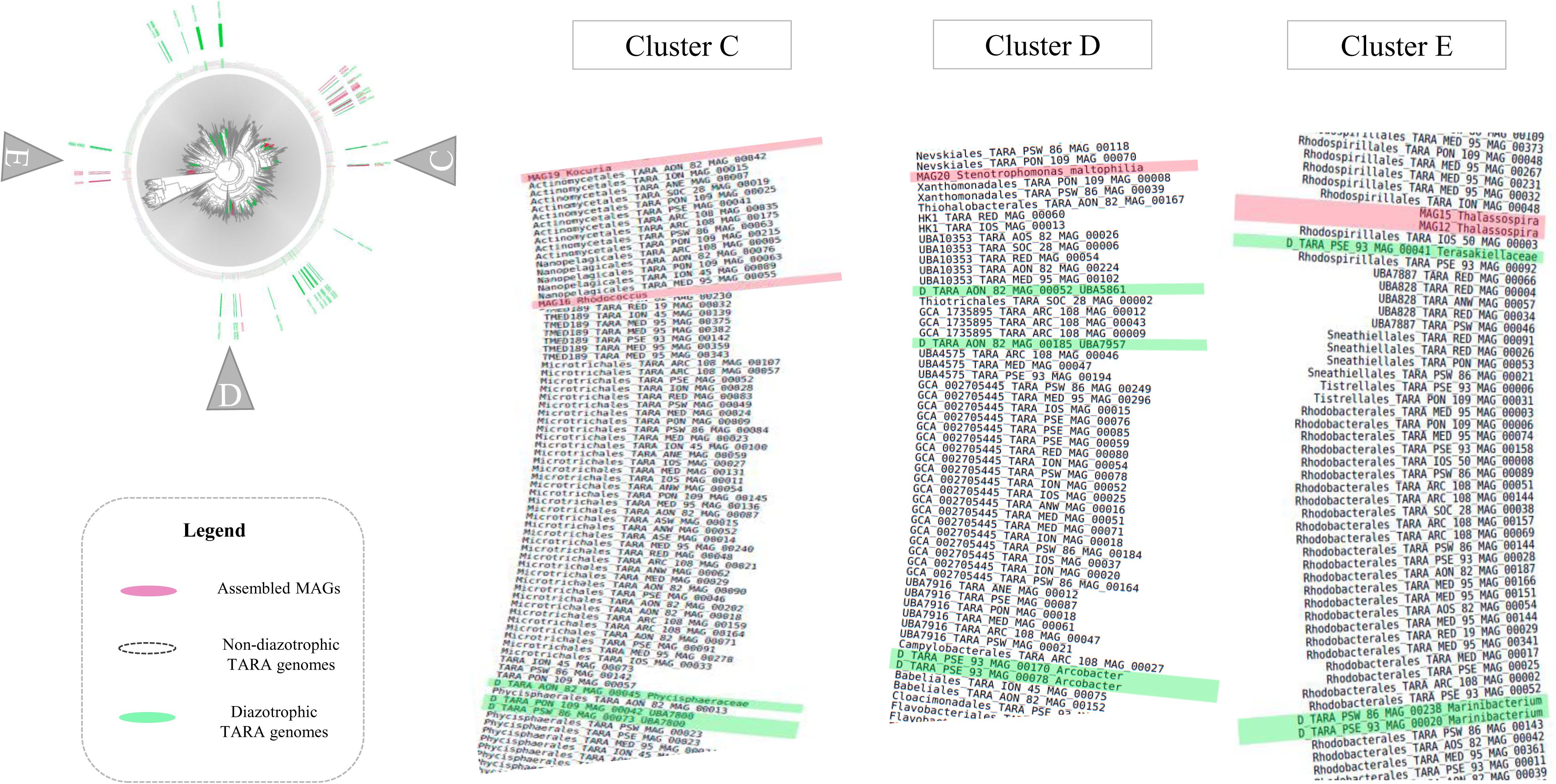
Phylogenomic tree representing clusters C, D and E MAGs from *Pt15* and other microalgae with 1888 TARA ocean bacterial MAGs.

**Figure S4.**
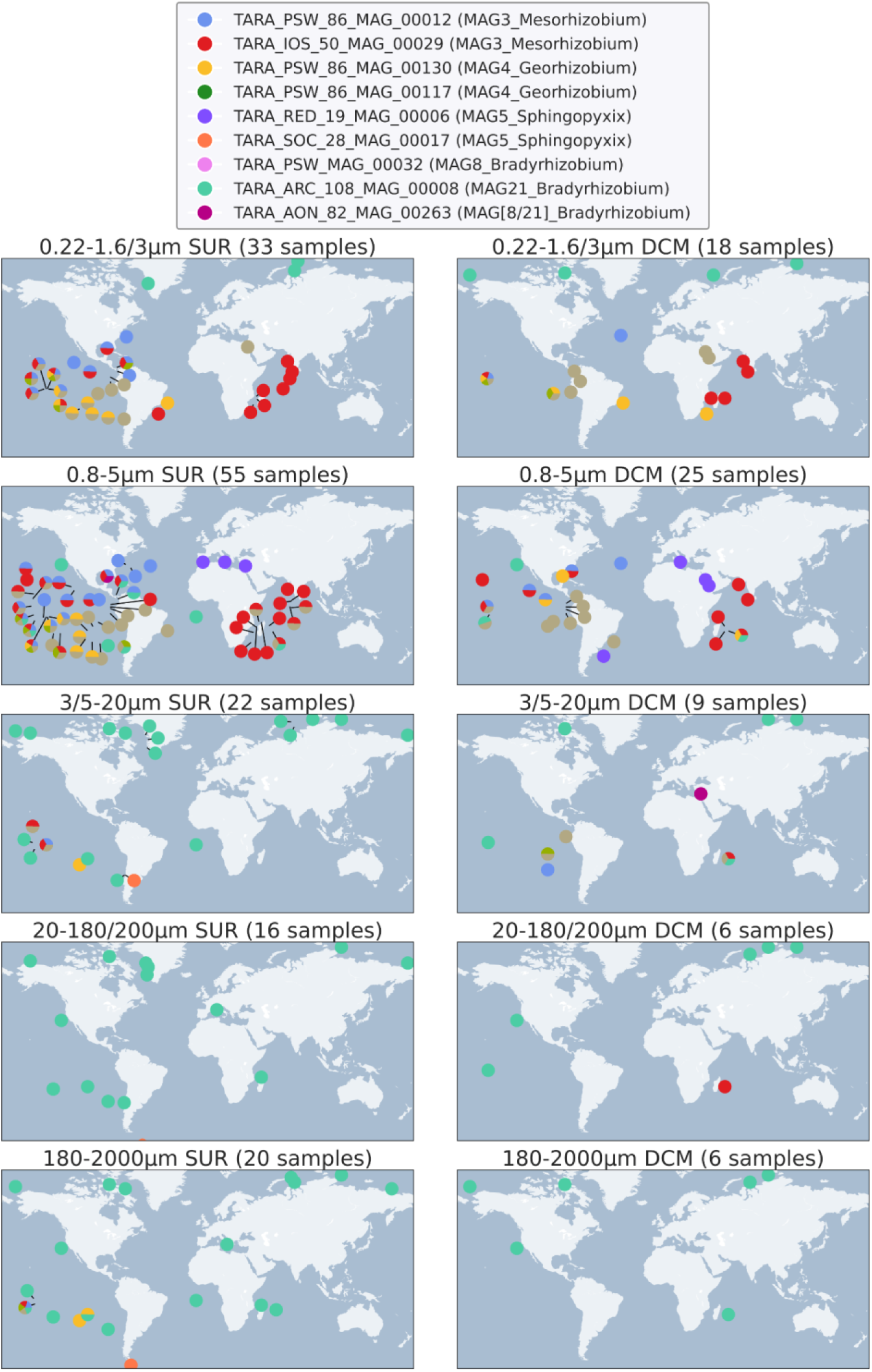
Geographical distribution of phylogenetically closest *Pt15* associated TARA oceans MAGs in different size fractions from surface and DCM samples.

**Figure S5.**
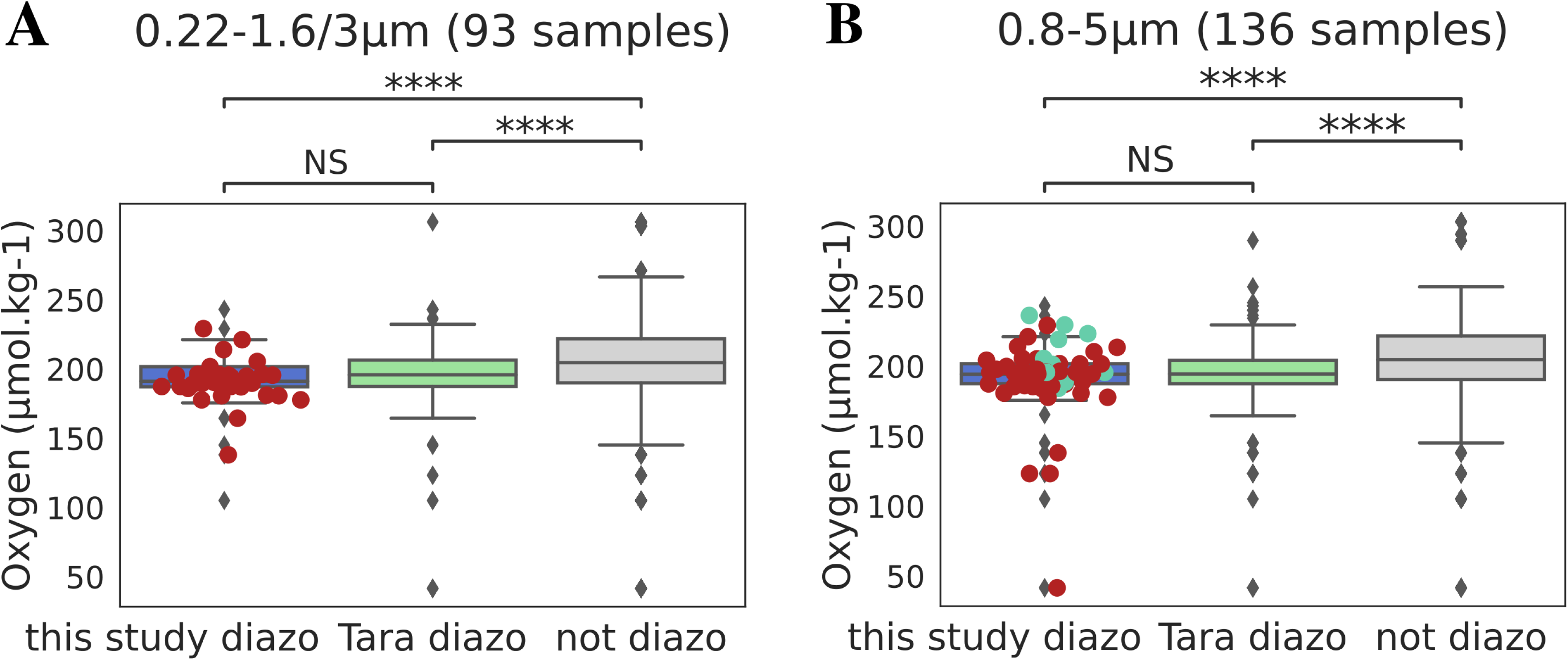
Relative abundance of the top 2 most occurring MAGs of *Pt15* bacterial community (this study diazo, *Mesorhizobium* MAG3 and *Bradyrhizobium* MAG8) compared to TARA diazotrophic MAGs and non-diazotrophic MAGs in the environment in respect to oxygen in 0.22-1.6/3 µm (A) and 0.8-5 µm (B) size fractions.

**Additional File 1.** Media recipes used for isolation of bacteria from *Phaeodactylum tricornutum* and other microalgae.

**Table S1.** Combined table representing amount of ethylene (Acetylene reduction assay), N15 incorporation assay and PON detection in xenic *Pt15* in nitrate deplete condition.

**Table S2.** Details of taxonomy for bacterial metagenomics data in nitrate deplete and nitrate replete conditions (replicates).

**Table S3.** MAG features, species-specific plasmid detection from *Pt15* meta-genomic data and TARA MAGs corresponding to Pt15 MAGs.

**Table S4.** Gene annotation using anvio_COG20, anvio_KEGG and anvio_Prokka of the metagenomics data in nitrate deplete and replete conditions (replicates).

**Table S5.** Details of Microalgae collection screened in nitrate deplete condition and bacteria isolated from specific microalgae.

**Table S6.** Features of the genomes selected from Mircoscope for phylogenetic studies.

**Table S7.** List of primers used in the study.

## Notes

### Competing Interest Statement

The authors have declared no competing interest.

### Summary of Updates

Clarifications of some results Figures with higher resolution

